# Distinct Strategies for Engineering Bioenergy Traits in Sorghum Drive Divergent Functional Shifts in the Rhizosphere Microbiome

**DOI:** 10.64898/2026.10.08.757756

**Authors:** Yen Ning Chai, Jutta Dalton, Yang Tian, Rita Kuo, Christopher De Ben, Daniel H. Putnam, Suzanne Kosina, Monikaben Nimavat, Edward Baidoo, Thomas Harwood, Katherine Louie, Benjamin Bowen, Christa Pennacchio, Robert Riley, Trent R. Northen, Aymerick Eudes, Corinne D. Scown, Henrik V. Scheller

**Affiliations:** Joint BioEnergy Institute, Environmental Genomics and Systems Biology Division, Lawrence Berkeley National Laboratory, Berkeley, CA 94720; Joint BioEnergy Institute, Biological Systems and Engineering Division, Lawrence Berkeley National Laboratory, Berkeley, CA, 94720; University of California-Davis, Davis, CA 95616; Joint Genome Institute, Lawrence Berkeley National Laboratory, Berkeley, CA 94720

**Keywords:** *Sorghum bicolor*, rhizosphere microbiome, metagenome-assembled genomes, TonB-dependent transporter, Carbohydrate-Active enZyme, plant cell wall degradation, deficit irrigation

## Abstract

Engineering bioenergy crops for enhanced biomass digestibility and high-value co-products is a promising approach to sustainable biofuel production. However, the impact of these engineered crops on the belowground microbiome remains poorly understood. In this study, we characterized the composition and functional responses of the belowground microbiome associated with transgenic sorghum lines: one with reduced cell wall crosslinking (AT10) and another with elevated protocatechuic acid (PCA) production (Qsub). The lines were grown under field conditions with four irrigation levels, and samples were collected twice during the growing season. We found that the two transgenic types drove a subtle but significant divergence in bacterial communities, both compositionally and functionally. Community-wide functional profiling showed that AT10 rhizospheres were consistently depleted in genes associated with mobile genetic elements, while Qsub rhizospheres were enriched in TonB-dependent transport and vanillate monooxygenase, an enzyme that funnels lignin-derived monomers toward PCA degradation, alongside reduced pectin- and cellulose-degradation genes. Genome-resolved analysis linked these community-level shifts to specific taxa: in Qsub, the enriched MAGs were taxonomically diverse, ranging from genome-reduced Patescibacteria with an episymbiotic lifestyle to taxa enriched in lignin-associated auxiliary activity enzymes. In AT10, the depleted Azospirillaceae MAGs carried unusually high mobile genetic element content, consistent with the community-wide pattern. Collectively, these findings offer new insights into how genetic modifications aimed at improving bioenergy traits shapes the composition and function of the rhizosphere microbiome, with potential downstream consequences for soil carbon cycling.

## Introduction

Overcoming cell wall recalcitrance in bioenergy feedstocks has remained a key focus in lignocellulosic biofuel and chemical production. While lignin is a major contributor to cell wall recalcitrance, it is not the sole factor that limits feedstocks degradability [1]. The level of feruloylation in grass cell walls is negatively correlated to cell wall digestibility [1–5]. Approximately 3% of grass cell wall dry weight is composed of ferulate (FA) esterified to arabinoxylans [4, 6, 7]. Arabinosyl FA can form covalent bonds with lignin and with each other, cross-linking the cell wall matrix, which significantly increases cell wall rigidity and hinders enzymatic degradation [8].

One promising strategy to reduce ferulate-mediated cross-linking of the cell wall matrix is to increase the level of *p*-coumarylation on arabinoxylans, as *p*-coumarate (pCA), which is also esterified to arabinoxylans, likely competes with FA for attachment sites. In our recent work [9], we successfully reduced cell wall recalcitrance in a sorghum line (hereafter referred to as AT10) by enhancing *p*-coumarylation on arabinoxylans through the overexpression of a rice acyltransferase (OsAT10). Sorghum is an increasingly important bioenergy crop, not only because it is high-yielding and familiar to farmers, but also because its genetic diversity [10] and phenotypic versatility [11] allow some varieties to thrive with reduced water and nutrient inputs. While reducing cell wall recalcitrance can lower the cost and resource requirements of producing biofuel from sorghum and other lignocellulosic feedstocks, the impacts of such cell wall modifications on soil microbial communities and carbon cycling are not well understood.

Lignin, as a major contributor to cell wall recalcitrance, is a polymer of phenolic building blocks. During cell wall degradation, lignin-derived phenolics are broken down by microbes into a few key intermediates, including protocatechuic acid (PCA), which are precursors to central carbon metabolism [12]. Well-known lignin-derived aromatic degraders include *Sphingobium sp.* SYK-6 [13], *Pseudomonas putida* [14], and *Rhodococcus* strains [15], all of which are of interest for the industrial valorization of lignin-derived aromatics. To understand the ecological implication of cell wall modification on soil ecosystem functioning, it is crucial to investigate how PCA influences soil microbial communities. As part of this study, we incorporated a sorghum line (hereafter referred to as Qsub) that accumulates PCA in biomass [16]. The accumulation of PCA in biomass and root exudate can serve as a proxy to understand the mechanistic impact of increased phenolic input on soil microbiome composition and functioning.

In addition to reducing cell wall recalcitrance, engineering plants to produce high-value bioproducts is a promising strategy to valorize feedstocks. PCA has high industrial value [17], making it a valuable bioproduct on top of biofuel production. Engineering feedstocks toward elevated phenolic production raises the flux of these compounds into soil. Evidence suggests that increased input of phenolic compounds from both plant litter and root exudates [18–20] can alter the composition and function of the soil or rhizosphere communities [21–25]. Phenolic compounds are major constituents of SOM and their effect on decomposition is context-dependent: while some phenolics can suppress microbial activity by inhibiting extracellular enzymes [27], simple aromatics such as 4-hydroxybenzoic acid and benzoic acid are readily metabolized and can stimulate soil microbial communities to decompose SOM more rapidly, enhancing carbon turnover and CO₂ emissions [21, 22, 28]. Therefore, the objective of this study was to assess the impact of not only reduced cell wall recalcitrance but also increased phenolic input on soil ecological processes, particularly carbon cycling, and to identify the microbial taxa involved in these processes.

In this study, we investigated the microbial communities associated with engineered sorghum lines grown under field conditions. Four irrigation levels corresponding to 100%, 75%, 50%, and 25% of sorghum evapotranspiration were included to evaluate whether transgenic lines differed from wild-type controls in agronomic performance under water deficit, and to survey whether the genotype-induced shifts in the rhizosphere microbiome were detectable across this gradient. We employed next-generation amplicon and metagenomic sequencing to assess changes in community composition and functional potential, respectively. We found that bacterial, but not fungal, communities in the rhizosphere were significantly impacted by the two transgenic types in an irrigation-dependent manner. As our primary interest was in genotype-driven rather than irrigation-driven changes to the rhizosphere microbiome, we restricted genome-resolved metagenomic sequencing to the full-irrigation treatment to characterize functional shifts induced by the transgenic lines. Metagenomic analyses suggested that the two transgenic lines also influenced the microbial function, with Qsub-enriched bacteria encoding CAZymes consistent with efficient scavenging and coupled breakdown-uptake of plant-derived polymers. The AT10 lines showed comparatively modest and less functionally coherent responses. Together, these findings offer new insights into how different strategies for engineering bioenergy traits can differentially impact soil ecological processes.

## Materials and Methods

### Field site

The field site was located in Davis, California, at GPS coordinates 38.53502 N, 121.77985 W. The soil type in this field is clay loam. The field was left fallow for the past two years and triticale was planted in winter 2021 as cover crop. The planting took place on June 6, 2022, in single row plots that measured 7.3 m x 0.76 m.

Nitrogen fertilizer was applied in two split applications: 123 kg N ha⁻¹ before planting on May 25 and an additional 39 kg N ha⁻¹ at the flowering stage on August 15. To control aphids, imidacloprid (40.4%) was applied twice at a rate of 12 oz acre⁻¹, on July 28 and August 15.

Plots were irrigated using a surface drip irrigation system. Four irrigation treatments corresponding to 25% (10.64 in), 50% (13.64 in), 75% (15.64 in), and 100% (23.64 in) of crop evapotranspiration (ET) were imposed, with drought treatments initiated on July 13, 2022.

Two transgenic sorghum types were tested, namely AT10 [9] and Qsub [29]. Three independent transgenic lines and a wild-type segregant were planted for each of these sorghum types, along with their background wild-type Wheatland line. A complete randomized block design was used, consisting of four blocks or replicates. Each block was divided into four sections, with each section randomly assigned to one of the four irrigation treatments. Within each section, nine sorghum genotypes were randomly assigned. Details about sampling and sample processing are presented in the Supplementary Materials and Methods.

### Sampling

We sampled the belowground microbiome twice during the growing season, at the sorghum boot stage (Aug 9, 2022) and the soft dough stage (Oct 12, 2022). These growth stages were selected because the root system is more established and the exudate-driven rhizosphere microbiome is expected to be more developed. Cumulative phenolic exudation has also been shown to increase over time in *Arabidopsis [30]*. During the microbiome sampling, bulk soil, rhizosphere, and roots (hereafter referred to as sample type) were sampled from each plot to a depth of approximately 25 cm.

Sorghum biomass was harvested Oct 11 and 13, for grain heads and stover, respectively. Details about the sampling procedure and sample processing are presented in the Supplementary Materials and Methods.

### Soil moisture and carbon analysis

Gravimetric soil moisture was measured by drying 30 g of fresh soil in a 65°C oven until a constant weight was reached, and the mass loss was recorded. Total and water-dissolvable organic carbon and nitrogen were measured using a multi N/C 2100S duo equipped with a CLD nitrogen detector and HT1300 solids analyzer (Analytik Jena, Jena, Germany). Details about soil organic carbon and nitrogen analyses are available in the Supplementary Materials and Methods.

### Extraction and analysis of root cell wall–bound phenolics and root exudates

Using the same root tissues utilized for microbiome profiling, we extracted cell wall-bound phenolics: pCA and FA for the AT10 lines, and PCA and PCA-glucoside for the Qsub lines. Only samples from the 25% and 100% irrigation treatments were included in this analysis. Details about the isolation of these phenolics are available in the Supplementary Materials and Methods.

### Statistical analysis

Non-microbiome data including biomass, soil chemistry, metabolite data were analyzed using the same approach with R Studio version 2025.05.0+496 [31]. An analysis of variance (ANOVA) was used to assess the effects of irrigation treatment, sorghum line, and their interaction, while accounting for block effect. Significant results were followed up by pairwise comparisons using Tukey’s honestly significant difference (HSD) test implemented in the *emmeans* package version 2.0.3 [32]. Compact letter displays were then generated from the Tukey-adjusted comparisons using ‘cld’ function in the *multcompView* package version 0.1-11 [33].

### Amplicon sequencing and data analysis

The V4 region of the bacterial 16S rRNA gene and Internal Transcribed Spacer 1 (ITS1) located between the fungal 18S and 5.8S rRNA genes were targeted to profile the bacterial and fungal communities, respectively. The primers used to amplify the V4 region of 16S rRNA genes were 515F (5’-GTGCCAGCMGCCGCGGTAA-3’) and 806R (5’-GGACTACHVGGGTWTCTAAT-3’) [34]; for ITS1 we used ITS1-1F-F (5’-CTTGGTCATTTAGAGGAAGTAA-3’) [35] and ITS1-1F-R (5’-GCTGCGTTCTTCATCGATGC-3’) [36]. Peptide nucleic acid (PNA) blockers were included during PCR amplification to reduce host chloroplast (plastid) and mitochondrial 16S rRNA amplification [37]. Amplicons from soil and rhizosphere samples were sequenced by Novogene Co., Ltd. on a NovaSeq platform (Illumina, San Diego, CA), and root samples were sequenced at our in-house sequencing facility on a MiSeq platform (Illumina, San Diego, CA). Only root samples from full irrigation and lowest deficit irrigation (25% irrigation) plots were submitted for ITS sequencing. Sequence preprocessing was conducted using USEARCH, and taxonomic assignments for 16S and ITS amplicons were performed using the SILVA and UNITE databases, respectively. Alpha-, beta-diversity analyses, PERMANOVA, and differential abundance analyses were conducted in R. Detailed amplicon data processing and statistical analyses are provided in the Supplementary Materials and Methods.

### Metagenomics sequencing and data analysis

We submitted 36 rhizosphere DNA samples used for amplicon sequencing to the Joint Genome Institute (JGI) for metagenomic sequencing. These were the rhizosphere samples associated with Qsub and AT10 lines that received 100% irrigation. Metagenomic libraries were sequenced using 2×150 nucleotide reads on the Illumina Novaseq platform. Raw reads were then subjected to preprocessing, error curation, assembly, binning, function annotation, and taxonomy assignment through the standard DOE JGI Metagenomic Workflow [38]. Metagenome-assembled genomes (MAGs) were reconstructed to identify genes of the bacteria enriched in the rhizosphere of transgenic lines. Detailed procedures for metagenomic assembly, binning, annotation, and downstream analyses are provided in the Supplementary Materials and Methods.

### Data availability

The 16s and ITS amplicon data are available at NCBI under the bioproject number PRJNA1205755. The metagenomic data are available at the JGI Data Portal (https://data.jgi.doe.gov) under the proposal number 510400.

## Results

### Biomass, soil, cell wall composition, and root exudate data

Sorghum aboveground biomass was machine-harvested at the soft-dough stage. The AT10 line had grain yield comparable to that of the control group, which consisted of the Wheatland wild type and the wild-type segregants of the Qsub and AT10 lines. In contrast, grain yield was significantly lower in Qsub lines (Fig. 1A). No significant difference in stover yield was observed between the transgenic lines and control (Fig. 1A). Deficit irrigations reduced grain yield but had no significant impact on stover biomass. The yield reductions in grain were similar between the 50% and 25% irrigations (Fig. 1B). Transgenic lines of both types exhibited similar yield reductions to the control across different deficit irrigation levels (Fig. S1A).

**Fig. 1.**
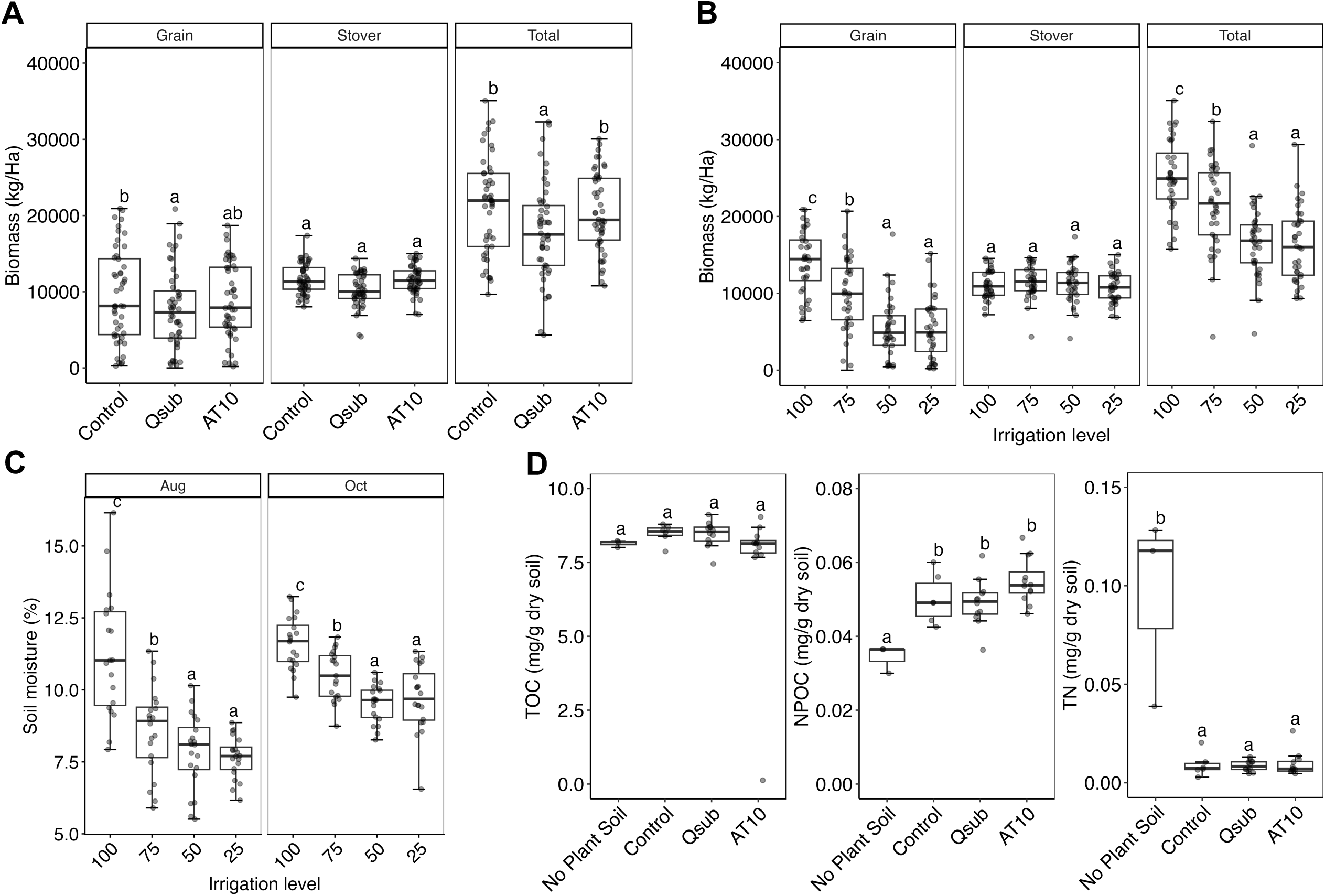
Plant biomass and soil characteristics. Dry yield of sorghum grain, stover, and total aboveground biomass by (**A**) sorghum line and (**B**) irrigation level. (**C**) Soil moisture content under each irrigation level. (D) Total organic carbon (TOC), water-dissolvable non-purgeable organic carbon (NPOC), and total nitrogen (TN) concentrations in the bulk soil. Different letters indicate significant differences based on Tukey’s HSD test (*p* < 0.05). Boxes represent the interquartile range with the median indicated by a horizontal line, and whiskers extend to values within 1.5× the interquartile range.

Soil moisture was measured and the deficit irrigation effect was greater during the August sampling compared with the October sampling (Fig. 1C). The soil moisture was similar between 50% and 25% irrigation treatments at both sampling time points, which was also reflected in the grain yield.

Soil total organic carbon (TOC), water-dissolvable non-purgeable organic carbon (NPOC), and total nitrogen (TN) were measured in August. NPOC was significantly higher in plant-associated soils compared with unplanted soil, whereas TN was substantially greater in unplanted soil (Fig. 1D). TOC did not differ significantly between plant-associated and unplanted soils (Fig. 1D), and neither the sorghum type nor the deficit irrigation treatment had a significant impact on TOC, NPOC, or TN (Fig. S1B).

In AT10 roots, we quantified FA- and pCA-arabinose levels and observed a significant increase in pCA-arabinose, along with a marginal decrease in FA-arabinose (Fig. S2A,B). The ratio of pCA- to FA-arabinose was also significantly greater in the AT10 lines compared with the control (Fig. S2C).

For Qsub lines, we measured PCA in roots collected from the field. PCA concentrations were markedly higher in the root of Qsub lines compared with the control (Fig. S2D). A hydroponic experiment was performed to measure the PCA in root exudates of Qsub lines. The amount of free PCA in the root exudate was comparable between Qsub lines and the control whereas the levels of PCA-glucoside were elevated in the Qsub lines (Fig. S2E). Both PCA and PCA-glucoside were highly abundant in the hydroponic root tissues of Qsub lines compared with the control (Fig. S2F).

### Compositional changes in microbial communities

16S and ITS amplicon sequencing were carried out to characterize the bacterial and fungal communities in soil, rhizosphere, and root endosphere (hereafter referred to as sample type) of these sorghum lines. Sample type was the dominant factor driving compositional changes in both bacterial and fungal communities, explaining 50.2% and 30.3% of the variation, respectively (*p* < 0.001) (Figs. 2 and S3). Harvest time, which reflects plant growth stage, exerted a stronger influence on the rhizosphere and root endosphere, accounting for an average of 5.4% of variation in bacterial communities and 8.9% in fungal communities (*p* < 0.001; Figs. 2A and S3A). In the soil, by contrast, harvest time accounted for 2.9% and 6.4% of variation in bacterial and fungal communities, respectively (*p* < 0.001; Figs. 2A and S3A). The October harvest showed an increase in Proteobacteria and Bacteroidota, along with a decrease in Actinobacteriota and Acidobacteriota compared with August (Fig. 2B). These changes were more pronounced in the root endosphere and rhizosphere than in the soil. For fungal communities, Ascomycota and Basidiomycota decreased, while Fungi_phy_incertae_sedis were enriched in the root endosphere relative to soil and rhizosphere (Fig. S3B). Both bacterial and fungal alpha diversity (Shannon entropy) was lowest in the root endosphere, intermediate in the rhizosphere, and highest in bulk soil and NPS (Fig. 2C, Fig. S3C). Bacterial diversity decreased significantly from August to October in both the rhizosphere and root endosphere (Fig. 2C). In contrast, fungal diversity increased in the rhizosphere, while the root endosphere remained unaffected by harvest time (Fig. S3C).

**Fig. 2.**
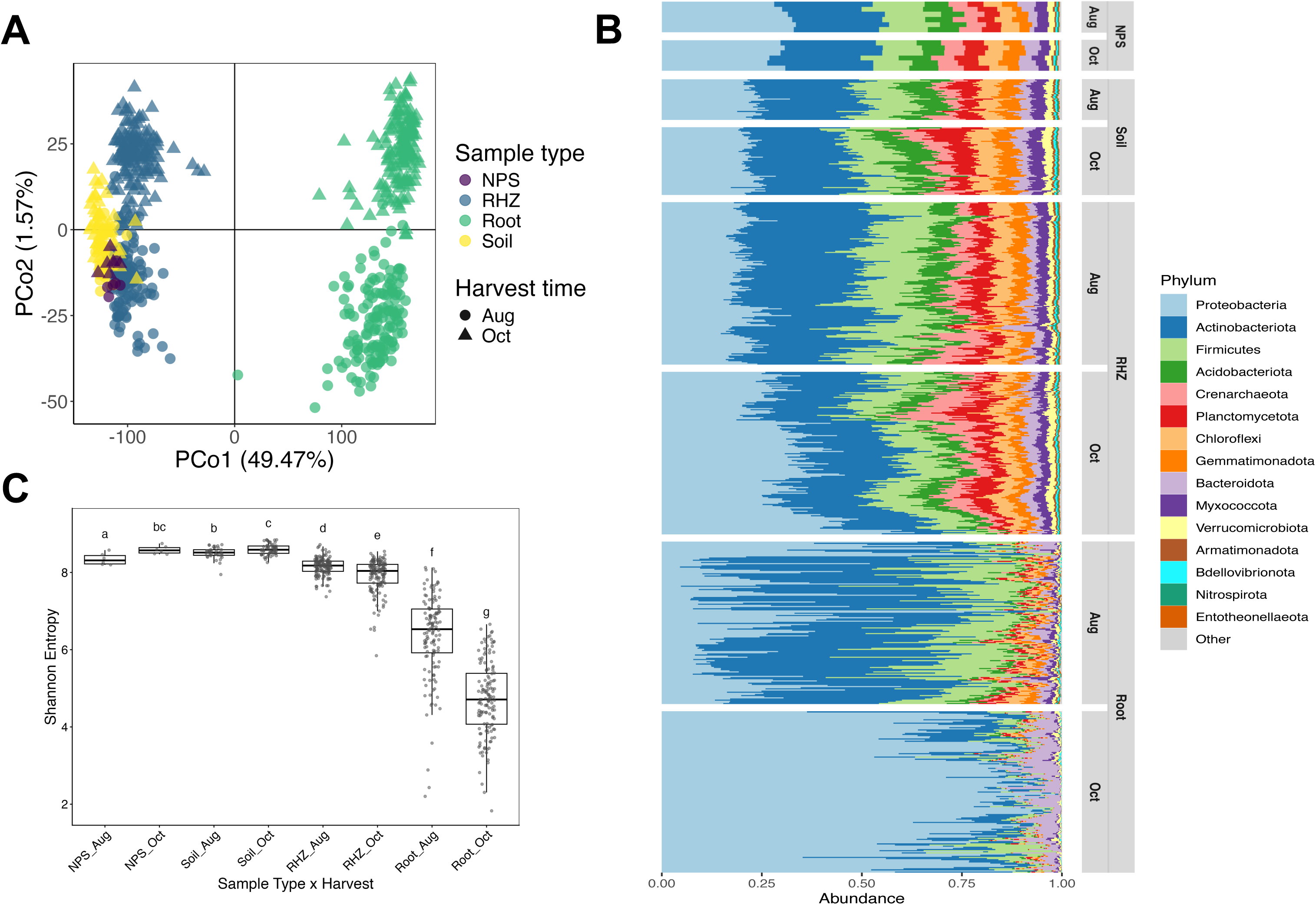
Bacterial community composition and alpha diversity across sample types and harvest time points. (**A**) Principal coordinates analysis (PCoA) based on Aitchison distance comparing bacterial communities across sample type and harvest time point. (**B**) Stacked bar plot showing the relative abundance of the 15 most abundant phyla grouped by sample type and harvest time point. **(C)** Shannon alpha diversity for each sample type and harvest time point combination.

Given the strong influence of sample type and harvest time, subsequent analyses were conducted separately for each sample type to assess the effect from deficit irrigation and sorghum line. Across sample types, deficit irrigation had the weakest effect on the root endosphere, where it accounted for 1.1% of variation in bacterial communities and 1.7% in fungal communities (*p* = 0.011 and *p* = 0.008, respectively), compared with the soil and rhizosphere, where it explained an average of 2.4% and 3.3% (*p* < 0.001), respectively. Despite these relatively modest effect sizes, deficit irrigation consistently altered the rhizosphere microbiome. Pairwise comparisons showed that every irrigation treatment differed significantly from one another for both bacterial and fungal communities (Supplementary table 1), in contrast to soil moisture and grain yield, where the 25% and 50% irrigation treatments did not differ significantly. In the root endosphere, bacterial community composition differed primarily between the 100% and 25% irrigation treatments, whereas fungal community composition was not significantly affected by irrigation (Supplementary table 1).

During the August harvest, we found a significant interaction between deficit irrigation and sorghum line in shaping rhizosphere bacterial communities (Fig. 3A), suggesting that the impact of sorghum line varied with water availability. Pairwise comparisons revealed that the Qsub rhizosphere community differed significantly from the control only under the 100% irrigation treatment, whereas the AT10 rhizosphere community differed significantly only under the 75% irrigation treatment (Fig. 3B, Table 1), with an average effect size of 4.45%. This interaction effect between deficit irrigation and sorghum line on the rhizosphere bacterial communities was only significant during the August harvest.

**Fig. 3.**
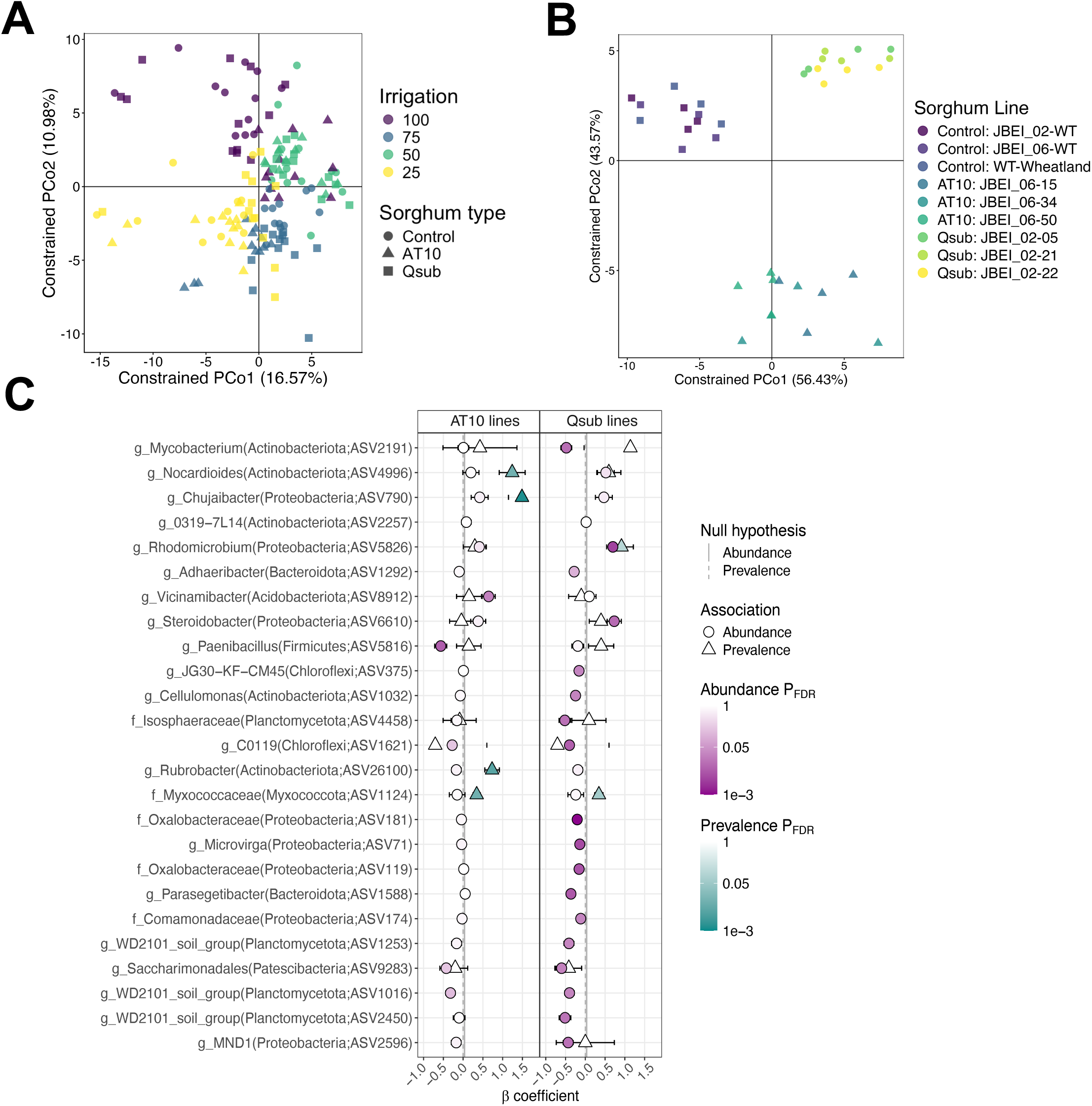
Impacts of deficit irrigation and sorghum line on rhizosphere bacterial communities. (**A**) Constrained analysis of principal coordinates (CAP) based on Aitchison distance showing the interactive effect of deficit irrigation and sorghum line. (**B**) Effect of sorghum line on bacterial composition under 100% and 25% irrigation. (**C**) Amplicon sequence variants (ASVs) showing differential abundance or prevalence in AT10 and Qsub lines relative to controls, as determined by MaAsLin3. In abundance models, coefficients represent log₂ fold changes in relative abundance, whereas in prevalence models, coefficients represent changes in the log-odds of ASV presence.

**Table1.** Pairwise PERMANOVA comparing the rhizosphere bacterial communities of AT10 and Qsub lines against the control (wild type and wild-type segregants) at each deficit irrigation level during the August harvest.

|  | R2 | F | P | Padj |
| --- | --- | --- | --- | --- |
| 100% |  |  |  |  |
| AT10 vs Control | 0.045 | 1.045 | 0.046 * | 0.110 |
| Qsub vs Control | 0.051 | 1.182 | < 0.001 *** | 0.002 ** |
| 75% |  |  |  |  |
| AT10 vs Control | 0.044 | 1.023 | 0.001 *** | 0.007 ** |
| Qsub vs Control | 0.039 | 0.895 | 0.196 | 0.335 |
| 50% |  |  |  |  |
| AT10 vs Control | 0.040 | 0.916 | 0.417 | 0.500 |
| Qsub vs Control | 0.039 | 0.885 | 0.389 | 0.500 |
| 25% |  |  |  |  |
| AT10 vs Control | 0.046 | 1.053 | 0.036 * | 0.107 |
| Qsub vs Control | 0.045 | 1.045 | 0.016 * | 0.062 |

In the AT10 rhizosphere, we found that *Rubrobacter* and *Nocardioides* (Actinobacteriota) and Mycococcaceae (Myxococcota) were more prevalent in the rhizosphere, indicating that they were detected in a greater proportion of samples. (Fig 3C). The Qsub rhizosphere was enriched in*Steroidobacter* and *Rhodomicrobium*, both belonging to the phylum Pseudomonadota (formerly Proteobacteria) (Fig 3C). Conversely, members of the order Saccharimonadales (Patescibacteria) and the WD2101 soil group (Planctomycetota) were depleted in the Qsub rhizosphere.

No significant effects of sorghum line were detected in the root endosphere bacterial community or in the fungal communities of either the rhizosphere or root endosphere.

### Community-level functional profiles are shaped by transgenic lines

To elucidate the functional aspects of the changes induced by transgenic sorghum lines, we conducted metagenomic sequencing on 36 rhizosphere DNA samples. These were the same samples used for amplicon sequencing, collected during August/boot stage and received full-irrigation. The metagenomic sequencing yielded 57.93 Gb of scaffolds (mean 1.61 Gb per sample) with an N50 of 1,048 bp and an average scaffold length of 1,022 bp. Across the 36 samples, an average of 53% of quality-filtered reads (ranging from 35.1% to 65.1%) were successfully assembled into contigs. The resulting assemblies had an average sequencing depth of 15.36x.

We investigated the functional shifts in the rhizosphere microbiome by performing a differential abundance analysis on KEGG Orthologs (KO) using Fisher’s Exact test. To pinpoint the core response to each transgenic type, we focused on genes that were consistently enriched or depleted across all three lines of AT10 and Qsub. In the AT10 lines, we detected a consistent depletion of numerous KOs related to mobile genetic elements (Fig 4A). For the Qsub line, we initially hypothesized an enrichment of genes encoding for PCA *meta*- and *ortho*-cleavage pathways, but their abundances were comparable to those in the control group (Fig 4B). Genes encoding the PCA *para*-cleavage pathway were not detected. Instead, we found a significant enrichment of genes encoding vanillate monooxygenase (vanAB), an enzyme located upstream that funnels lignin monomers into the PCA pathway (Fig 4C). The most significant change we observed was an enrichment of several TonB transporter components, although their abundance was only moderately higher than that of the control (Fig 4D). We also observed an enrichment of genes encoding the xylobiose transport system, along with reductions in genes encoding pectin transporter (alpha-1,4-digalacturonate permease) and lytic cellulose monooxygenase across all Qsub lines (Fig 4C).

**Fig. 4.**
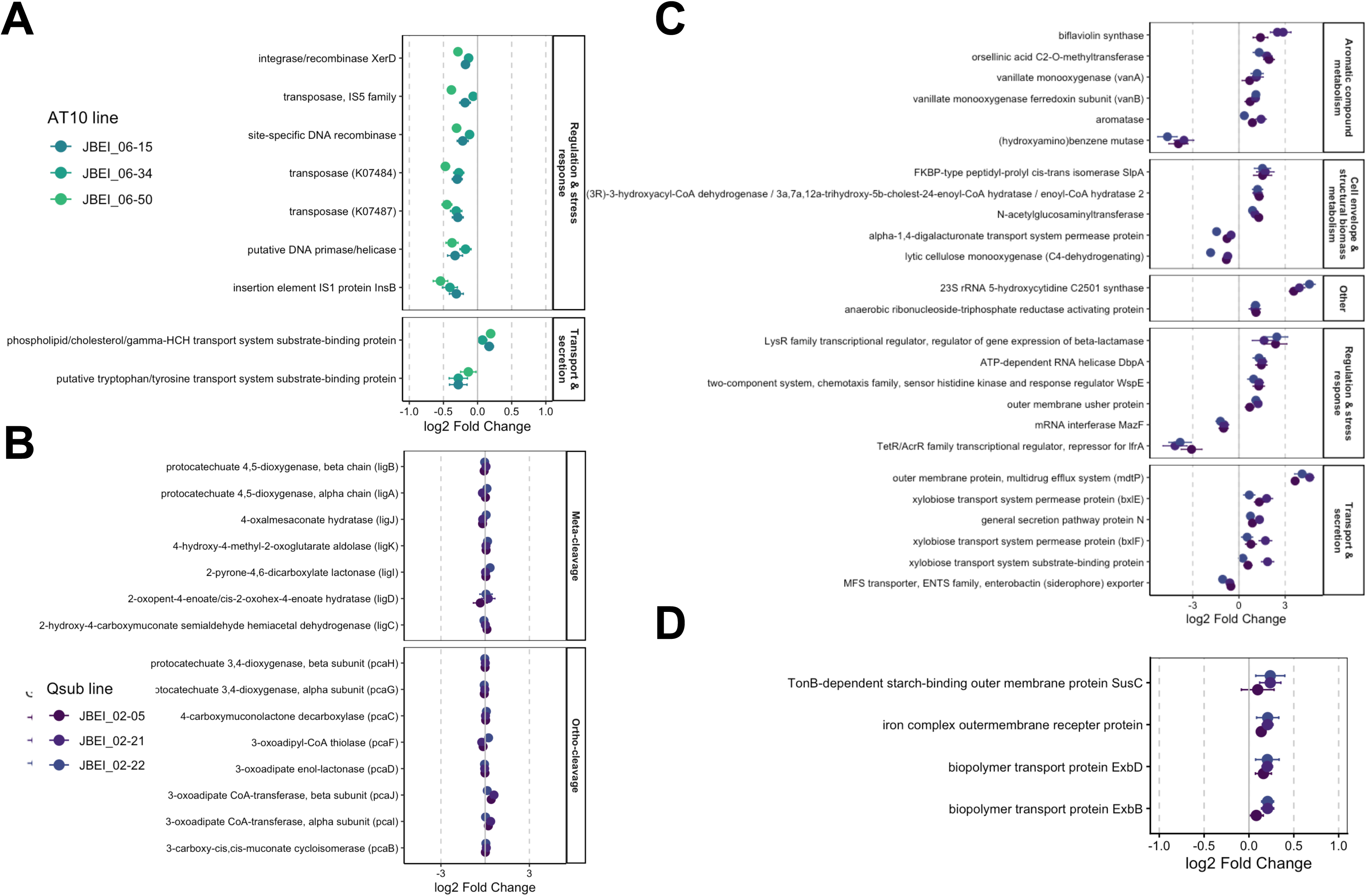
Differential gene abundance in the sorghum rhizosphere metagenome. (**A**) KEGG Orthology (KO) terms differentially enriched or depleted in the AT10 rhizosphere compared with controls. (**B**) Genes encoding the protocatechuate (PCA) degradation pathways in the Qsub lines. (**C**) KO terms differentially enriched or depleted in the Qsub rhizosphere compared with controls. (**D**) Genes encoding TonB transporter components in the Qsub rhizosphere. Statistical analysis was performed using Fisher’s exact test with FDR adjustment for multiple comparisons.

### Genome-resolved metagenomics identifies microbial taxa associated with AT10 and Qsub lines

We then assembled microbial genomes from the metagenomic data to gain insight into the ecological roles of microbes influenced by our transgenic lines. The 36 rhizosphere samples were assembled both individually and collectively: individual-sample assembly to reduce complexity and yield high-quality contigs, and co-assembly to enable cross-sample comparison and detect low-abundance genomes. This combination of approaches yielded a total of 205 medium-to high-quality unique MAGs (completeness ≥50%; contamination ≤10%; 200 bacteria and 5 archaea; Supplementary Table 2) after dereplication at the species level (>95% ANI). Of these, 155 MAGs were recovered from the single-sample assemblies and 34 from the co-assembly, and 11 were detected in both, collectively spanning 18 bacterial phyla and one archaeal phylum (Fig 5).

**Fig. 5.**
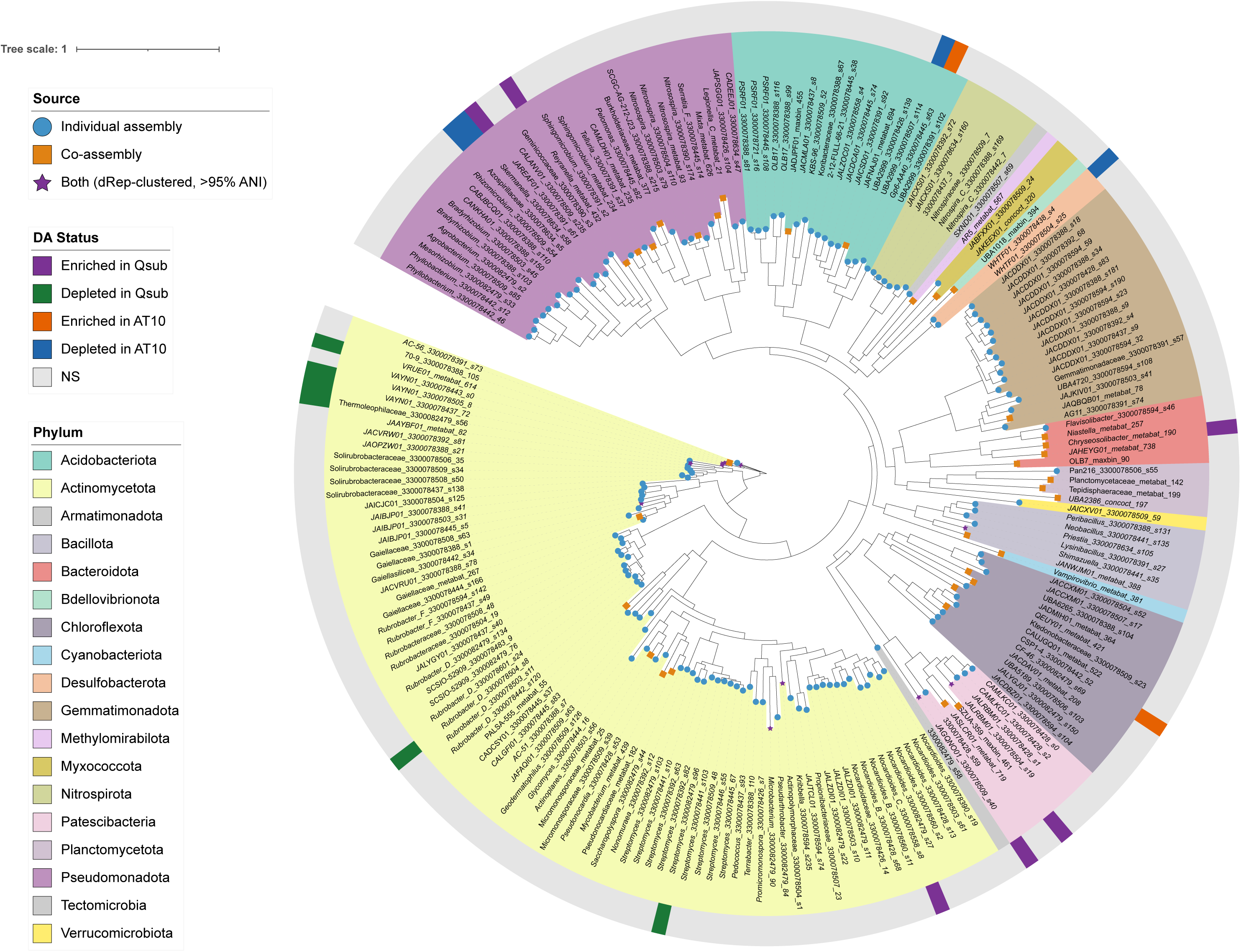
Phylogeny of MAGs. (**A**) **Phylogenetic tree of 200 medium-to-high quality bacterial metagenome-assembled genomes (MAGs);** five archaeal MAGs were excluded from the tree. The outer ring denotess differentially abundant MAGs in the AT10 and Qsub lines. Tip shapes indicate whether each MAG was recovered from individual assembly, co-assembly, or both.

To identify MAGs affected by the transgenic lines, we performed a differential abundance analysis using DESeq2. Eighteen out of 205 MAGs were differentially enriched or depleted relative to the control (Fig. S4). In the AT10 lines, MAGs belonging to family Azospirillaceae, including the genus *Skermanella*, were depleted, and this was consistent with the 16S results. These Azospirillaceae MAGs also carried substantially more mobile genetic elements (41 and 35 genes) than the other two AT10-enriched MAGs, CSP1-4 (2 genes) and Gp6-AA40 (3 genes).

Most of the differentially abundant MAGs (12 of 18) were associated with the Qsub line. Strikingly, all six depleted MAGs belonged to the phylum Actinobacteriota. Four belonged to the order *Solirubrobacterales* (families 70-9 and *Thermoleophiliaceae*), whereas the remaining two belonged to the genera *Rubrobacter_D* and *Streptomyces*. In contrast, the MAGs enriched in the Qsub rhizosphere spanned multiple bacterial phyla, including Actinobacteriota (*Nocardioides_C*), Patescibacteria (*Saccharimonadales*), Bacteroidota (*Chryseosolibacter*), and Alphaproteobacteria (*Reyranella* and JAREAF01).

Notably, one Solirubrobacterales MAG (3300078505_8) and one Saccharimonadales MAG (3300078509_s40), both of which were recovered in the individual and co-assemblies, were identified as differentially abundant only in the co-assembly analysis. These MAGs were abundant and consistently detected across samples in the co-assembly, whereas in the individual assemblies they were recovered from only one or two samples and were therefore excluded from the differential abundance analysis because they did not meet the prevalence threshold.

### Differentially abundant MAGs exhibit distinct plant cell wall degradation potential

We next analyzed the Carbohydrate-Active enZyme (CAZyme) profiles of the differentially abundant MAGs to determine if they differed in capacity in degrading plant cell wall (Fig 6A). We focused on CAZymes involved in the degradation of major plant cell wall components and grouped them based on the substrate: cellulose, hemicellulose, pectin, and lignin.

**Fig. 6.**
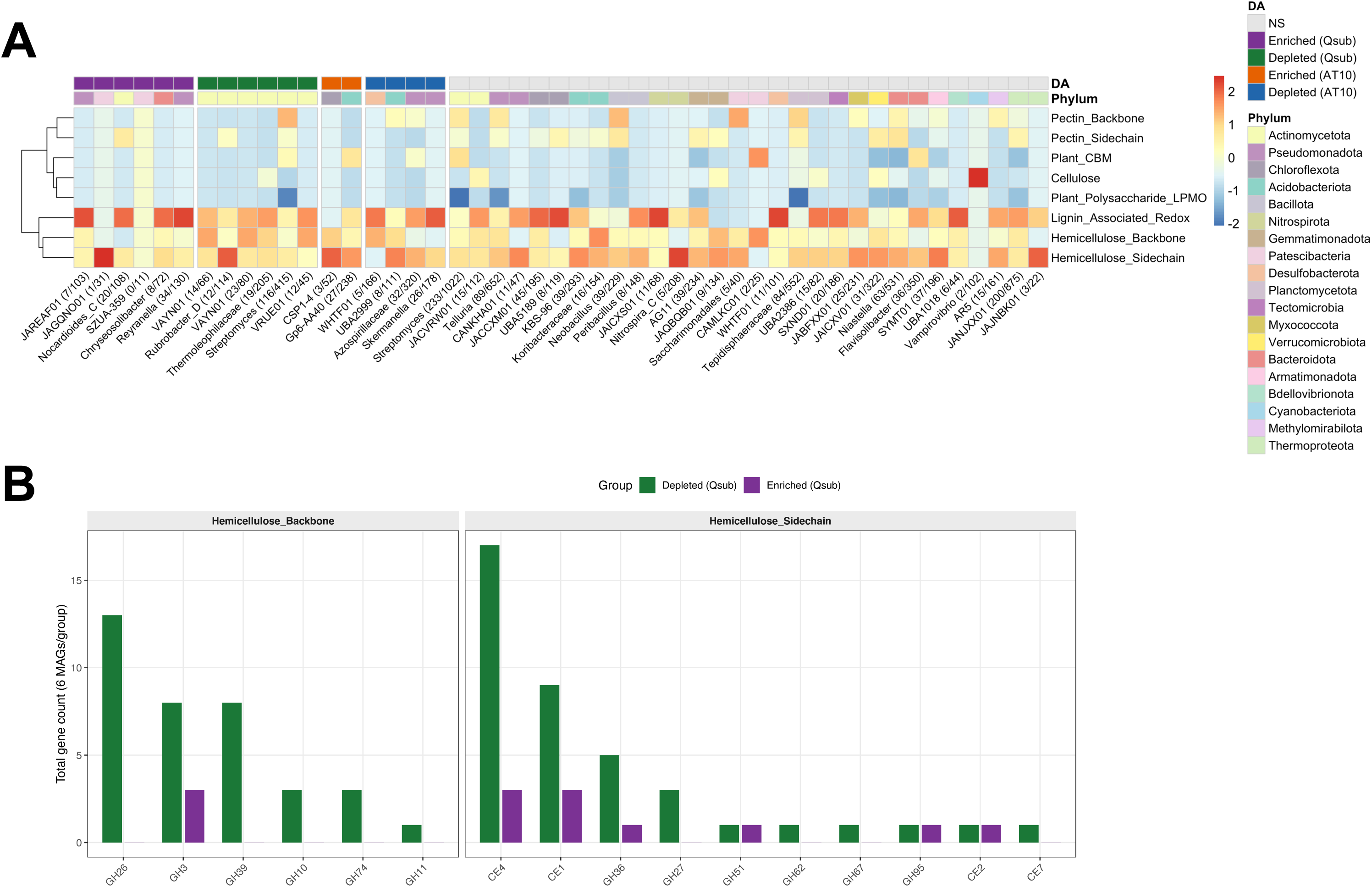
Carbohydrate-active enzyme (CAZyme) profiles of differentially abundant MAGs. (**A**) Heatmap of column-scaled, log₁₀-transformed CAZyme category abundance in the 18 differentially abundant MAGs and representative non-significant MAGs from each phylum.Values in parentheses after MAG IDs denote the proportion of plant cell wall degradation CAZymes relative to the total identified CAZyme repertoire. **(B)** Abundance of hemicellulose degrading CAZymes in Qsub-enriched and Qsub-depleted MAGs. CAZyme categories include cellulose degradation (GH6, GH7, GH9, GH45, GH48); hemicellulose backbone degradation (GH3, GH10, GH11, GH26, GH39, GH74); hemicellulose side-chain degradation (GH27, GH36, GH51, GH54, GH62, GH67, GH95, GH115, CE1, CE2, CE4, CE5, CE6, CE7, CE15, CE16); pectin backbone degradation (GH28, PL1, PL3, PL4, PL9, PL10, PL11); pectin side-chain degradation (CE8, CE12, GH53, GH78, GH127); plant polysaccharide-active lytic polysaccharide monooxygenases (AA9, AA14); lignin-associated auxiliary activity enzymes (AA1, AA2, AA3, AA4, AA6); and plant cell wall-binding carbohydrate-binding modules (CBM1, CBM2, CBM3, CBM6, CBM13). Abbreviations: GH: glycoside hydrolase; PL: polysaccharide lyase; CE: carbohydrate esterase; AA: auxiliary activities.

In the AT10 rhizosphere, the small number of differentially abundant MAGs (n = 2 enriched, n = 4 depleted) limited our ability to identify meaningful trends in CAZyme composition. In Qsub, the depleted MAGs encoded a significantly greater number of hemicellulose backbone-degrading CAZymes than the enriched ones (DESeq2, Padj < 0.05; Fig 6B). This pattern was largely driven by the four Solirubrobacterales MAGs, which have approximately 40% on average of hemicellulose backbone-degrading CAZymes out of the total number of plant cell wall degrading CAZymes. The remaining depleted MAGs, *Streptomyces* and *Rubrobacter_D*, encoded broader repertoires of plant cell wall-degrading CAZymes. Notably, the *Streptomyces* MAG had the largest genome among the differentially abundant MAGs in Qsub and encoded a number of CAZymes involved in the degradation of cellulose, hemicellulose, pectin, and lignin. *Rubrobacter_D* also encoded CAZymes targeting all of these substrates except cellulose.

Among the MAGs enriched in Qsub rhizosphere, two Patescibacteria MAGs encoded little or no plant cell wall-degrading CAZymes and had markedly smaller genomes than the other differentially abundant MAGs (approximately 0.8 Mbp versus 3 Mbp) (Supplementary table 2). The remaining enriched MAGs encoded proportionally more lignin-associated auxiliary activity enzymes (AA1, AA3, AA4, and AA6), which accounted for an average of 73% of their annotated plant cell wall-related CAZymes, compared with 34% in the depleted MAGs.

## Discussion

In this study, we have confirmed that modifying plant cell wall crosslinking and increasing PCA production in sorghum can influence the rhizosphere bacterial communities both compositionally and functionally. The differentially abundant taxa were largely different between AT10 and Qsub lines, suggesting that each line provides different resources and therefore selects for a different subset of soil microbial community. Qsub-enriched MAGs occupied diverse niches, ranging from genome-reduced taxa with minimal degradative capacity to lignin-redox specialists. In comparison, the effects of the AT10 line were more modest, with fewer differentially abundant taxa that do not show a clear trend in substrate specificity.

Although the AT10 line affected relatively few taxa, both depleted MAGs belonged to Azospirillaceae (one identified as the genus *Skermanella*), and both were also detected as differentially abundant taxa in the 16S dataset. Notably, both MAGs had relatively large genomes and encoded numerous mobile genetic elements [39]. This was consistent with the community-level reduction in mobile genetic elements observed in the AT10 rhizosphere. This convergence between community- and genome-level patterns suggests that AT10 selects against taxa with high genomic plasticity, potentially favoring a more genomically stable rhizosphere community.

In the Qsub line, several PCA-degradation genes showed no sign of enrichment: PCA 3,4- and 4,5-dioxygenase abundances were comparable to control, and neither PCA 2,3-dioxygenase nor PCA decarboxylase were detected in our dataset. This was consistent with the 16S data, which showed no enrichment of well-known PCA-degrading genera such as *Sphingomonas* [40], *Caulobacter* [41], and *Burkholderia* [42]. Instead, we found significant enrichment of vanillate monooxygenase, an enzyme acting upstream of PCA in the lignin-funneling pathway, together with several TonB-dependent transporter components. TonB is a tripartite, energy-dependent transport system restricted to Gram-negative bacteria that has been implicated in the uptake of large lignin-derived aromatics such as the biaryl 5,5′-dehydrodivanillate [43]. Direct involvement in PCA uptake seems unlikely, however, since PCA is typically transported by the far less energy-intensive PcaK permease [44]. TonB expression is nevertheless known to be induced by vanillate in *Pseudomonas aeruginosa* [45], raising the possibility that PCA and related phenolics such as vanillate serve as a cue for TonB-mediated capture of larger lignin-derived aromatics.

Genome-resolved metagenomics revealed that the Qsub line reshaped the functional composition of the rhizosphere microbiome along taxonomic lines. All depleted MAGs belonged to Actinobacteriota, but they varied in degradative capacity: *Streptomyces* and *Rubrobacter_D* encoded broad repertoires of plant cell wall-degrading CAZymes, whereas Solirubrobacterales were narrowly specialized in mannan degradation. The enriched MAGs, by contrast, spanned multiple phyla and functional strategies, including genome-streamlined Saccharimonadales with minimal plant cell wall-degrading capacity [50], Bacteroidota enriched in polysaccharide degradation genes [51], and taxa enriched in lignin-associated auxiliary activity enzymes. This pattern suggests that elevated PCA availability does not simply favor broad-spectrum biomass degraders, but instead restructures the community around a more diverse set of specialized carbon-acquisition strategies.

Our data also suggest that deficit irrigation restructures the rhizosphere microbiome by enriching for members of Actinobacteriota, which is consistent with another sorghum drought study [54]. However, the magnitude of this enrichment was less pronounced in our study, likely reflecting differences in how the deficit was imposed: our treatment extended the interval between irrigation events, whereas Xu et al. imposed a complete cessation of irrigation [54]. Notably, grain yield and biomass declines were comparable between the 25% and 50% deficit irrigation treatments, even though the two treatments produced distinct rhizosphere community structures, suggesting that the magnitude of plant stress may not scale directly with the extent of microbiome restructuring. Furthermore, the effects of the transgenic lines depended on water availability. The AT10 line significantly altered the rhizosphere bacterial community under the 75% irrigation treatment, whereas the Qsub line had its strongest effect under full irrigation. This genotype-by-irrigation interaction is consistent with a previous rice study showing that host genotype effects on the rhizosphere microbiome are drought dependent, likely because drought alters root metabolite profiles [55].

The functional shifts observed in the Qsub rhizosphere may have important implications for soil carbon cycling. Both Bacteroidota [56] and *Nocardioides* [57] are well known for their capacity to degrade plant cell wall polysaccharides. Their enrichment in the Qsub rhizosphere may therefore indicate the availability of accessible or favorable carbon substrates selected for these taxa. However, the ecological consequences of this shift remain uncertain. On one hand, such specialization could trigger a “priming effect” which occurs when simple compounds, such as phenolic acids, activate specialist soil microbes that co-metabolize native soil organic matter, resulting in a net loss of soil carbon stocks and elevated CO₂ efflux [21, 22]. This outcome would ultimately undermine soil carbon sequestration efforts. Conversely, if the enriched specialists exhibit high carbon use efficiency, they may preferentially channel plant-derived substrates into their own biomass. The resulting microbial necromass is a key precursor to new, stable mineral-associated organic matter, which would enhance long-term carbon storage. Therefore, the ultimate impact of this specialist-driven community on the soil carbon balance is ambiguous and likely depends on factors like microbial carbon use efficiency and substrate preference, which warrant further investigation.

The individual-assembly approach may fail to recover genuinely differentially abundant taxa. Because each individual-assembly MAG is built solely from one sample’s own contigs, reads from other samples often fail to map back to it. As a result, genuinely abundant taxa can appear nearly absent in every sample except the one they were assembled from, artificially suppressing their apparent abundance elsewhere. This explains why several MAGs identified as significant in the co-assembly analysis were not detected in the individual-assembly analysis. We also observed discrepancies between the 16S and MAG-based analyses. For example, Saccharimonadales was Qsub-depleted in the 16S dataset but enriched in the MAG analyses. Likewise, both *Rhodomicrobium* and *Steroidobacter* were significantly enriched in Qsub in the 16S dataset, but *Rhodomicrobium* had no corresponding MAG at all, likely reflecting naming differences between taxonomy databases (SILVA for 16S vs. GTDB for MAGs) or the higher taxonomic resolution of genome-based classification. *Steroidobacter*, while present in the metagenome, was not significantly enriched, possibly because the 16S and MAG data were capturing different species within this taxonomically diverse family (over 40 distinct Steroidobacteraceae ASVs were detected). In contrast, the depletion of Azospirillaceae in the AT10 rhizosphere was consistently identified by both methods. Because MAG-based analysis offers higher taxonomic and functional resolution, we prioritized the MAG results throughout this study. Overall, it is valuable to have both datasets available, as metagenomics provides functional insight while 16S provides complementary compositional information.

Our study provides a snapshot of the compositional and functional changes induced by altering cell wall traits and increasing the exudation of phenolics. While we found some interesting trends, this change could be transient, as the compositional change was most pronounced during the boot stage, and we only profiled the metagenome at that single timepoint due to resource constraints.

Although our metagenomics and 16S data provided complementary insights into the taxonomic and functional composition of the rhizosphere microbiome, these approaches capture gene potential rather than gene activity. Transcriptomic profiling would be a valuable next step to determine whether functional categories such as PCA degradation genes, though stable in abundance, are differentially expressed. Integrating gene expression data with the compositional and functional patterns identified here would help clarify the mechanistic link between microbial community structure and root exudate degradation in the rhizosphere.

Taken together, our findings demonstrate that distinct strategies for engineering bioenergy traits in sorghum, whether by altering cell wall crosslinking or by increasing phenolic acid production, elicit divergent and taxonomically distinct responses in the rhizosphere microbiome. Rather than uniformly suppressing or stimulating microbial activity, each engineering strategy appears to select for specific functional guilds suited to the altered chemical environment it creates, from mobile element loss in AT10 to a diversification of lignin- and phenolic-associated metabolic strategies in Qsub. These results underscore that the ecological consequences of crop engineering are trait-specific rather than generic, and they highlight the rhizosphere microbiome as an important, and currently underappreciated, factor to consider when designing bioenergy crops for both improved biomass processing and co-product valorization.

## Acknowledgements

We thank Hsiao-Han Lin, Karen Serrano, Lorenzo Washington, Minliang Yang, Sami Kazaz, Nick Downs, Sasilada Sirirungruang for volunteering in the samplings.

## Author contributions

YNC, AE, HVS, and DHP designed the study; CDB planted and maintained the study site; YNC, JD, YT, and HVS collected samples at the study site; RK performed amplicon sequencing; SK conducted soil carbon analyses and interpreted the results; MN, EB, TH, KL, BB, and CP performed and interpreted metabolomic analyses; YNC analyzed amplicon and metagenomic sequencing data; RR performed metagenomic co-assembly; CP coordinated the metagenomic sequencing; TN, CS, and HVS supervised the study; and YNC wrote the first draft of the manuscript, with all authors reviewing and editing the final version.

## Conflicts of interest

The authors declare no conflict of interest. T.R.N. is a founder of two non-profits, Prosper Soils and Bioaligned Labs with prior approval from LBNL.

## Funding

This work conducted at the Joint BioEnergy Institute was supported by the U.S. Department of Energy, Office of Science, Biological and Environmental Research Program, through contract DE-AC02-05CH11231 between Lawrence Berkeley National Laboratory and the U.S. Department of Energy. The work conducted by the U.S. Department of Energy Joint Genome Institute (https://ror.org/04xm1d337), a DOE Office of Science User Facility, was supported by the Office of Science of the U.S. Department of Energy operated under Contract No. DE-AC02-05CH11231.

## Supplementary Materials and Methods

### Sampling

We sampled the belowground microbiome twice during the growing season, around sorghum boot stage (Aug 9, 2022) and during soft dough stage (Oct 12, 2022). Bulk soil, rhizosphere, and root were sampled from each plot to a depth of approximately 25 cm. During the microbiome sampling, two plants were randomly excavated from each plot, one from each end of the rows, while preserving the middle 3.7 m for biomass measurements. Bulk soil was shaken off the plants and collected in ziplock bags. Rhizosphere was collected by vortexing representative subsamples of the roots from both plants in 30 mL of phosphate buffer (6.33 g/L NaH_2_PO_4_, 8.5 g/L Na_2_HPO_4_ anhydrous) in 50-mL tubes using a vortexer for one minute. After shaking, the root tissues were blotted dry and transferred to a new tube. Both the root and rhizosphere were kept on ice until they were brought back to the lab on the same day. Soil from unplanted edges and alleys was collected as controls.

For the biomass harvest, grain heads and stovers were harvested separately, on Oct 11, and 13, respectively. The grain heads were hand-harvested and weighed, 10 heads per plot were subsampled and dried for dry weight measurements. Stovers were machine-harvested from the middle 3.7 m of each plot for fresh weight, and approximately 1 kg of fresh stovers was subsampled for dry weight measurement. The grains and stovers were dried in an oven at 55 °C until they dried completely.

### Sample processing

In the lab, soil samples were sieved with a 0.5 mm sieve to remove root residues and aggregates. Part of the sieved soil was stored at -20 °C for microbial DNA extraction and the rest was used for soil moisture and soil organic matter analysis. The processing of rhizosphere and root samples was carried out as described (1). In brief, the mixture of rhizosphere and phosphate buffer was filtered using a 100 µm cell strainer. The flow-through was centrifuged, the supernatant was discarded, and the pelleted rhizosphere was resuspended in 2 mL of phosphate buffer, followed by transfer to a new 2 mL tube and storage at -20 °C for genomic DNA extraction. The root samples were surface-sterilized using 25% bleach, followed by 70% ethanol, and 3 water rinses for one minute each with shaking using a vortexer. Surface-sterilized roots were blotted dry, cut into small pieces using a sterilized pruner, and stored at -80 °C. The frozen root tissues were ground using a SPEX Sample Prep freezer mill model 6875D (SPEX Sample Prep, Metuchen, NJ). Genomic DNA from the soil and root samples was extracted using DNeasy PowerSoil Pro (Qiagen, Germantown, MD) following the manufacturer’s protocol. The DNA concentration was measured using Qubit and Synergy H1 Multi-Mode Reader with QuantiFluor dsDNA dye (Promega, Madison, WI) using 504 nm excitation and 531 nm emission wavelengths.

### Soil moisture and carbon analysis

For total organic carbon (TOC) analysis, 0.5 g of fresh soil was treated with 200 µL of 10% HCl to remove inorganic carbonates and dried in an oven at 105 °C. The remaining carbon was measured after combustion at 950 °C with a maximum integration time of 390 seconds, starting at 0.12 ppm with a threshold of 1.5 ppm and 10-count stability. Measurements were reported on a dry weight basis. For water-dissolvable non-purgeable organic carbon (NPOC) and total nitrogen (TN) analyses, 0.5 g of fresh soil was first extracted with 2 mL of HPLC-grade water and shaken overnight at 4°C. The soil slurry was centrifuged at 10,000 x g for 5 minutes at 10°C, and the supernatant was subsequently filtered through a 0.22 µm polyethersulfone filter. The filtrate was then diluted by adding 4 mL of HPLC-grade water to 1.5 mL of the filtrate. After thorough mixing, a 1.5 mL aliquot of the diluted solution was analyzed for NPOC and TN using a multi N/C 2100S duo equipped with a CLD nitrogen detector and HT1300 solids analyzer (Analytik Jena, Jena, Germany). Samples were processed using a 100 µL volume, with a 180 s purge (purge 1) followed by a 10 s purge (purge 2), an 800 °C furnace temperature, a 300-second integration time, a starting threshold of 0.12 ppm, and concentration thresholds of 0.2 ppm for carbon (C) and 0.5 ppm for nitrogen (N), all maintaining 7-count stability. Measurements were reported as parts per million (ppm) of water-dissolvable carbon or nitrogen.

### Extraction and analysis of root cell wall–bound phenolics and root exudates

To isolate xylan-bound pCA and FA, we first prepared cell wall residues (CWR) from 30 mg of lyophilized, ground root tissue. The tissue was washed three times with 80% ethanol (20 minutes each), followed by a 20-minute acetone wash, a 1:1 methanol:chloroform wash for 20 minutes, and a final 20-minute acetone wash; all steps were performed at room temperature. Samples were then dried overnight in a fume hood. To determine FA-arabinose and *p*-CA-arabinose, 10 mg of CWR was hydrolyzed in 50 mM trifluoroacetic acid (TFA) at 100 °C for two hours as previously described (2). After hydrolysis, the samples were centrifuged, and the TFA was removed from the supernatants by drying with a speed vacuum. The resulting pellet was resuspended in 500 µL of 50% methanol, filtered through a 3-kDa cut-off centrifugal filter, and diluted 40-fold for HPLC-ESI-QTOF-MS analysis using the method described previously (2). PCA-glucoside and PCA were extracted from Qsub roots using 80% methanol as described previously (3). Briefly, 30 mg of lyophilized, ground root tissue was extracted three times with 1 mL of 80% methanol at 70 °C. The extracts were pooled and combined with 1.5 mL of HPLC-grade water, centrifuged to remove debris, and passed through a 3-kDa centrifugal filter. Half of the sample was analyzed directly by HPLC-ESI-QTOF-MS to determine free PCA. The remaining half was subjected to acid hydrolysis and ethyl acetate partitioning to release the PCA aglycone, which was then analyzed by HPLC-ESI-QTOF-MS. A standard curve was generated using PCA standards ranging from 1.56 µM to 200 µM.

Because root exudates could not be collected during the field experiment, we conducted a complementary growth chamber experiment to verify that protocatechuic acid (PCA) is detectable in the root exudates of the Qsub lines. We grew two Qsub genotypes (JBEI 02-05, JBEI 02-22) and Wheatland wild type hydroponically in a growth chamber with a 16-h photoperiod. The chamber conditions were maintained at 25 °C and 80% relative humidity. Seeds were germinated in soil and seedlings were transferred to hydroponic containers containing full-strength Hoagland solution (4) with continuous aeration. Each line included five biological replicates, and root exudates were collected from a pool of five plants per replicate. Exudates were harvested at the 6-leaf stage. Prior to collection, the Hoagland solution was replaced with HPLC-grade water, and roots were incubated in it for 4 h. Shoot and root biomass were also recorded. Both root and exudate samples were immediately frozen at –80 °C and lyophilized for subsequent metabolomics analysis. Samples were submitted to the Joint Genome Institute (JGI) for LC-MS/MS analysis. PCA and PCA-glucosides were detected using a HILIC-Z column under negative ion mode using the method as described (5).

### Amplicon data analysis

For the 16S amplicons, demultiplexed raw reads were merged, primer-trimmed, quality checked (maximum expected error cutoff of 1 was used), and clustered into amplicon sequence variance (ASV) using USEARCH version 10.0.24 (6). The ITS amplicons underwent similar processes, except that *dada2* version 1.36.0 (7) was used instead of USEARCH, and primer sequences were retained. 16S and ITS taxonomy assignments were done using SILVA version 138.2 (8) and UNITE version 9 (9) database, using a Naive Bayesian Classifier (10,11). Subsequent statistical analyses and data visualizations were performed using RStudio version 2025.05.0+496 (12). *Phyloseq* version 1.52.0 (13) was used to organize the metadata, sequences, and ASV tables. For the 16s dataset, plant derived reads were removed using ‘subset_taxa’ function in *phyloseq* by excluding taxa assigned to mitochondria or chloroplasts. Host-derived reads accounted for an average of 0.08%, 0.11%, 0.30%, and 9.20% of reads in the NPS, bulk soil, rhizosphere, and root samples, respectively. After removal of host-derived reads, the 16S dataset contained a median of 160,622 reads per NPS sample (range: 129,703–174,698), 157,116 reads per bulk soil sample (range: 99,294–177,763), 158,184 reads per rhizosphere sample (range: 92,347–180,539), and 3,386 reads per root sample (range: 437–43,623). For the ITS dataset, the median sequencing depth was 98,814 reads per NPS sample (range: 94,723–181,194), 101,064 reads per rhizosphere sample (range: 94,909–190,737), 98,056 reads per rhizosphere sample (range: 79,149–174,200), and 25,518 reads per root sample (range: 4,897–110,283). For the alpha diversity analysis across all sample types, sequencing depth was rarefied to 1,000 reads per sample, and samples with less than 1,000 reads were dropped. Shannon diversity among groups were first assessed using the Kruskal-Wallis test, followed by pairwise Wilcoxon rank-sum tests with false discovery rate (FDR) adjustment for multiple comparisons.

For beta-diversity analysis, we used the Aitchison distance (Euclidean distance on centered-log-ratio [CLR]–transformed counts), which accounts for the compositional nature of sequencing data, rather than a relative-abundance–based dissimilarity such as Bray–Curtis or weighted UniFrac (14,15). We chose Aitchison because it measures communities through ratios between ASVs, which remain relatively stable when adding or dropping other taxa from the table. In contrast, Bray–Curtis and weighted UniFrac operate on relative abundances, so a change in one ASV alters the apparent abundances of unrelated ASVs, and they give greater weight to abundant taxa (16). Zeros were handled by adding a pseudocount of 1e-6 prior to the CLR-transformation, global dispersion patterns were computed using the ‘pcoa’ function in *ape* package version 5.8-1 (17) and visualized using *ggplot2* version 4.0.1 (18). Constrained Analysis of Principal Coordinates (CAP) was used to assess the effects of the experimental variables on the microbiome. Group dispersions or variances in the data were checked using the ‘betadisper’ function in *vegan* package version 2.7-2 (19). PERMANOVA was run using the ‘adonis2’ function to evaluate the effect size and statistics for each experimental variable. Because the sequencing depth was particularly uneven for root samples, we assessed whether the beta diversity results were consistent after accounting for depth. We tested for depth-community associations (correlation between library size and ordination axis scores), ran PERMANOVA models with sequencing depth included as a covariate, and confirmed that the results were similar regardless of depth.

Maaslin3 was used to identify the bacterial and fungal ASVs that were differentially affected the different experimental variables using both abundance and prevalence models (21). The abundance model tests whether the relative abundance of an ASV differs among experimental groups, whereas the prevalence model tests whether the probability of detecting an ASV (presence/absence) differs among groups. For the abundance analysis, a linear mixed-effects model was fitted with harvest stage, irrigation treatment, and sorghum line as fixed effects and block as a random effect. For the prevalence analysis, a logistic mixed-effects model with the same fixed and random effects was used to assess differences in ASV occurrence among experimental groups.

### Metagenomics sequencing and data analysis

Raw reads were curated using bbcms version 38.90 in *BBtool* package (<u>sourceforge.net/projects/bbmap/</u>). The assembly was performed separately for each sample and collectively for all samples. Individual sample assembly was conducted using metaSPAdes assembler version 3.15.2 (22) whereas co-assembly was done with MetaHipMer2 version 2.2.1.0 (23). Functional and taxonomic annotation was performed using IMG Annotation Pipeline (IMGAP v5.2.2). To identify genes enriched in the rhizosphere of transgenic lines, we used Fisher’s Exact Test with Benjamini-Hochberg (BH) adjustment to control for false discovery rate (FDR < 0.05). Comparisons were based on estimated gene copy number (KEGG Ortholog, KO, term annotations), calculated for each metagenome and compared between each transgenic line and its corresponding wild-type segregant. To identify robust, construct-wide effects, only KO terms that were significantly differentially abundant in all three independent transgenic lines within a given construct (Qsub or AT10) were retained.

For binning metagenome-assembled genomes (MAGs), MetaBat2 version 2:2.15 (24) and Semibin2 version 2.2.0 (25) were used for the individually assembled samples. For the co-assembly, CONCOCT version 1.1.0 (26), MaxBin version 2.2.7 (27), and MetaBat were used. Das Tool version 1.1.6 (28) was then used to integrate the bins generated by the co-assembly binning tools and remove redundancies by selecting the highest-quality bins. MAG quality was assessed using CheckM version 1.2.1 (29), only medium quality and above were retained (completeness > 50%, contamination <10%). We then pooled all the MAGs generated from both individual sample assembly and co-assembly, removing redundant MAGs using dRep version 3.6.2 (30) with an Average Nucleotide Identity (ANI) threshold of 95%. GTDB-tk version 2.1.1 was used for taxonomy assignment using database R207_v2 (31). CAZyme annotation of the MAGs’ genomes was conducted using Hidden Markov Models (HMMs) from the dbCAN2 CAZyme database implemented within KBase (32–34).

DESeq2 was used to identify differentially abundant metagenome-assembled genomes (MAGs). Prior to differential abundance analysis, low-abundance MAGs were filtered by retaining only those detected in at least two samples with a total count of at least five reads across all samples. After filtering, 78 of the 172 MAGs from the individual assemblies were retained, whereas all 45 MAGs from the co-assembly met the filtering criteria. Differential abundance analysis was performed to identify the MAGs enriched or depleted in each transgenic line while accounting for block effects. Each transgenic line was compared with its corresponding wild-type segregant and the Wheatland wild-type, and P values were adjusted for multiple testing using the Benjamini–Hochberg false discovery rate (FDR) method. IQ-TREE version 3.0.1 [31] was used to build a maximum-likelihood tree using the 200 bacterial MAG alignments produced in GTBTK, excluding the six archaeal MAGs, and the tree was visualized using iTOL version 7.2.2 [32].

## Notes

### Competing Interest Statement

The authors have declared no competing interest.

