## Supplemental Figures for "Distinct Strategies for Engineering Bioenergy Traits in Sorghum Drive Divergent Functional Shifts in the Rhizosphere Microbiome"

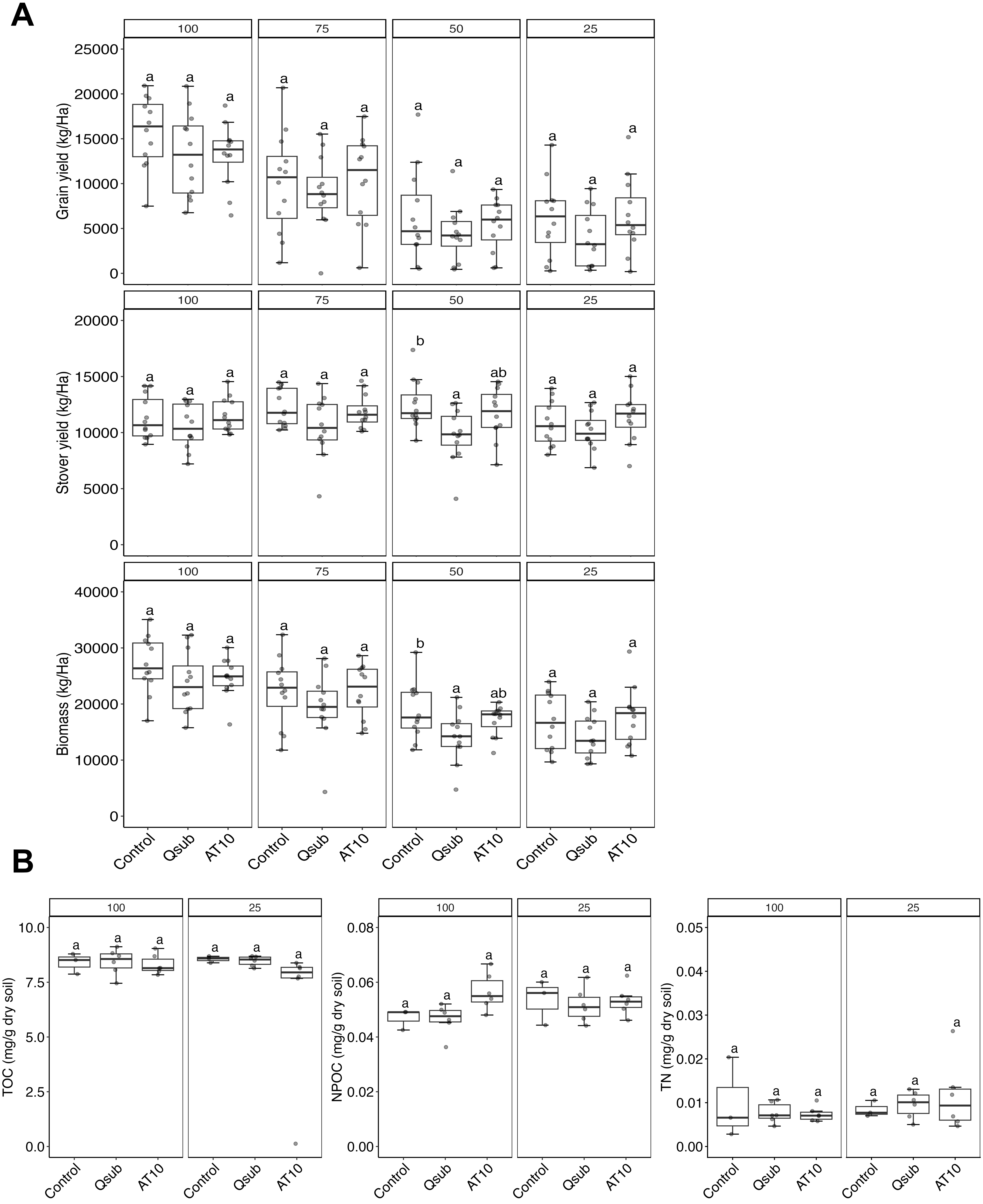


**Fig. S1. Interaction effects of sorghum line and irrigation level on (A) sorghum yield and (B) total organic carbon (TOC), non-purgeable organic carbon (NPOC), and total nitrogen (TN).** Different letters indicate significant differences based on Tukey's HSD test ($p < 0.05$).

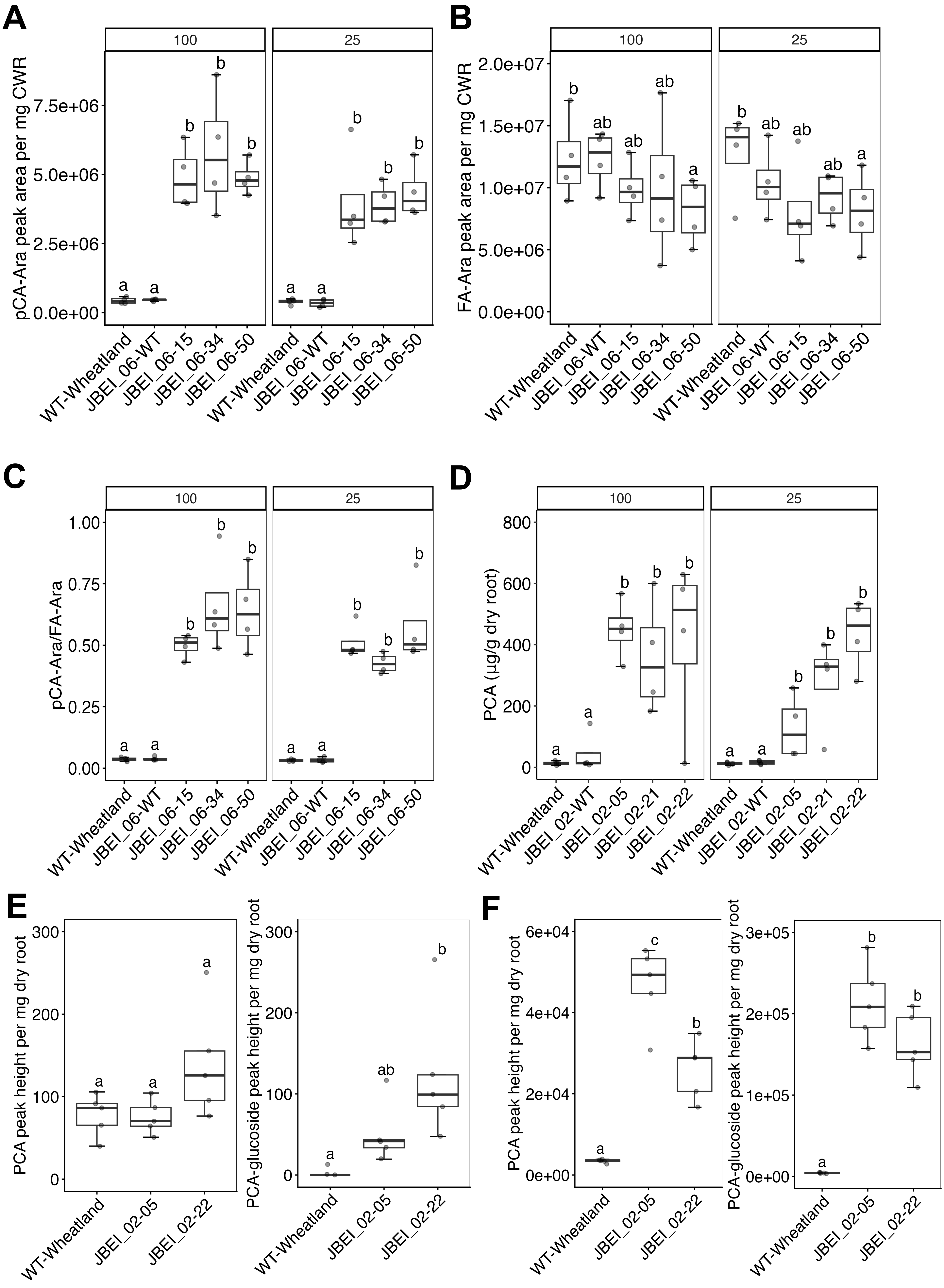
A
B
C
D
E
F
100
25
pCA-Ara peak area per mg CWR
FA-Ara peak area per mg CWR
pCA-Ara/FA-Ara
PCA (µg/g dry root)
PCA peak height per mg dry root
PCA-glucoside peak height per mg dry root
WT-Wheatland
JBEI_06-WT
JBEI_06-15
JBEI_06-34
JBEI_06-50
JBEI_02-WT
JBEI_02-05
JBEI_02-21
JBEI_02-22

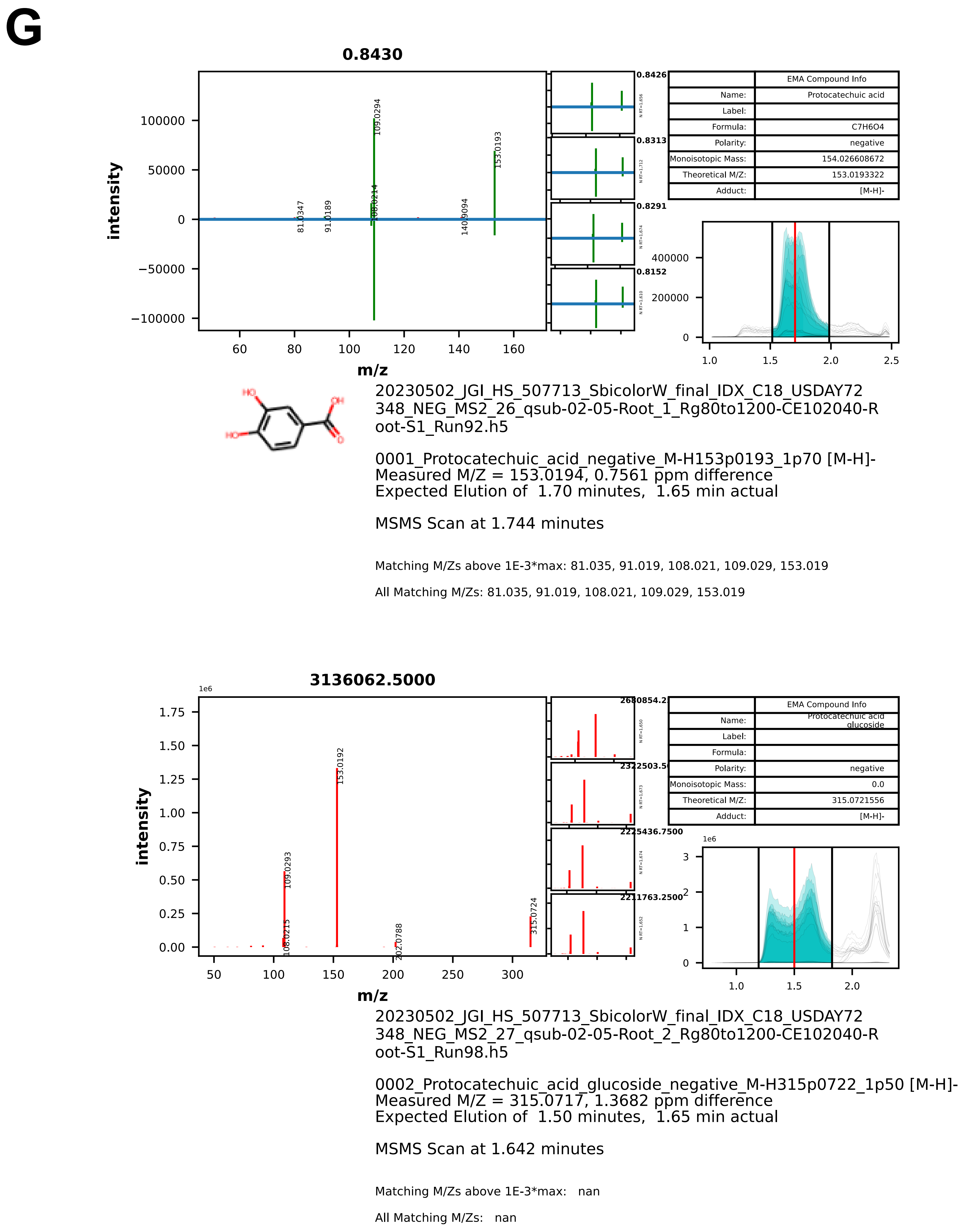


**Fig. S2. Phenolics in roots and root exudates.** (**A**) Content of *p*-coumaroyl-arabinose (pCA-Ara), (**B**) feruloyl-arabinose (FA-Ara), and (**C**) their ratio in the root cell wall residue (CWR) of AT10 and controls. (**D**) Protocatechuate (PCA) content in Qsub and control roots in field-grown plants. PCA and PCA glucoside concentrations in the (**E**) root exudates and (**F**) roots of hydroponically grown Qsub and control. (**G**) Extracted ion chromatograms and mirror plots of PCA (top) and PCA-glucoside (bottom). Different letters indicate significant differences based on Tukey's HSD test ($p < 0.05$).

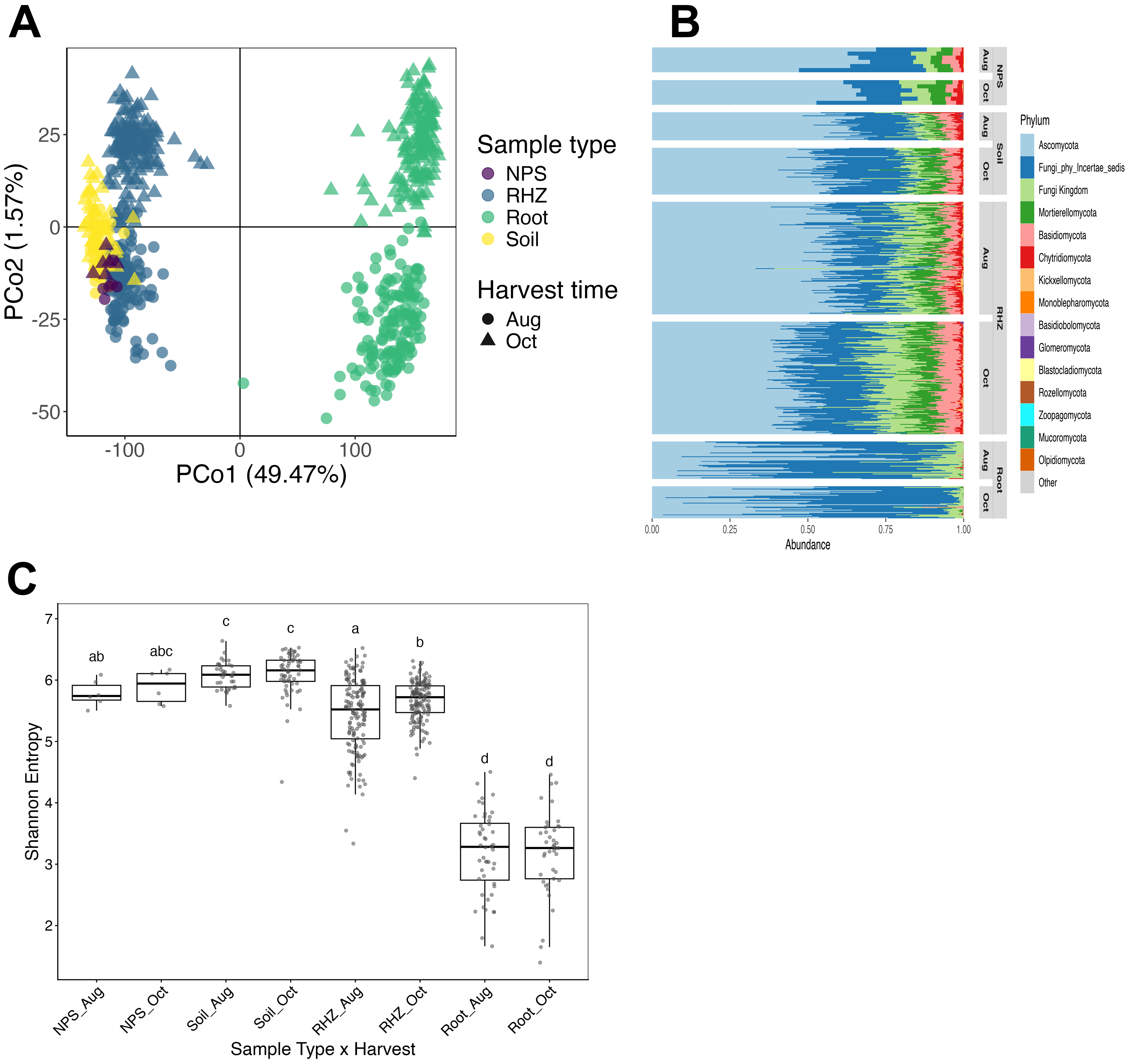


**Fig. S3. Fungal community composition and alpha diversity across sample types and harvest time points.** (**A**) Principal coordinates analysis (PCoA) based on Aitchison distance comparing fungal communities across sample types and harvest time points. (**B**) Stacked bar plot showing the relative abundance of the 15 most abundant phyla grouped by sample type and harvest time point. (C) Shannon alpha diversity for each sample type and harvest time point combination.

### Fig. S4

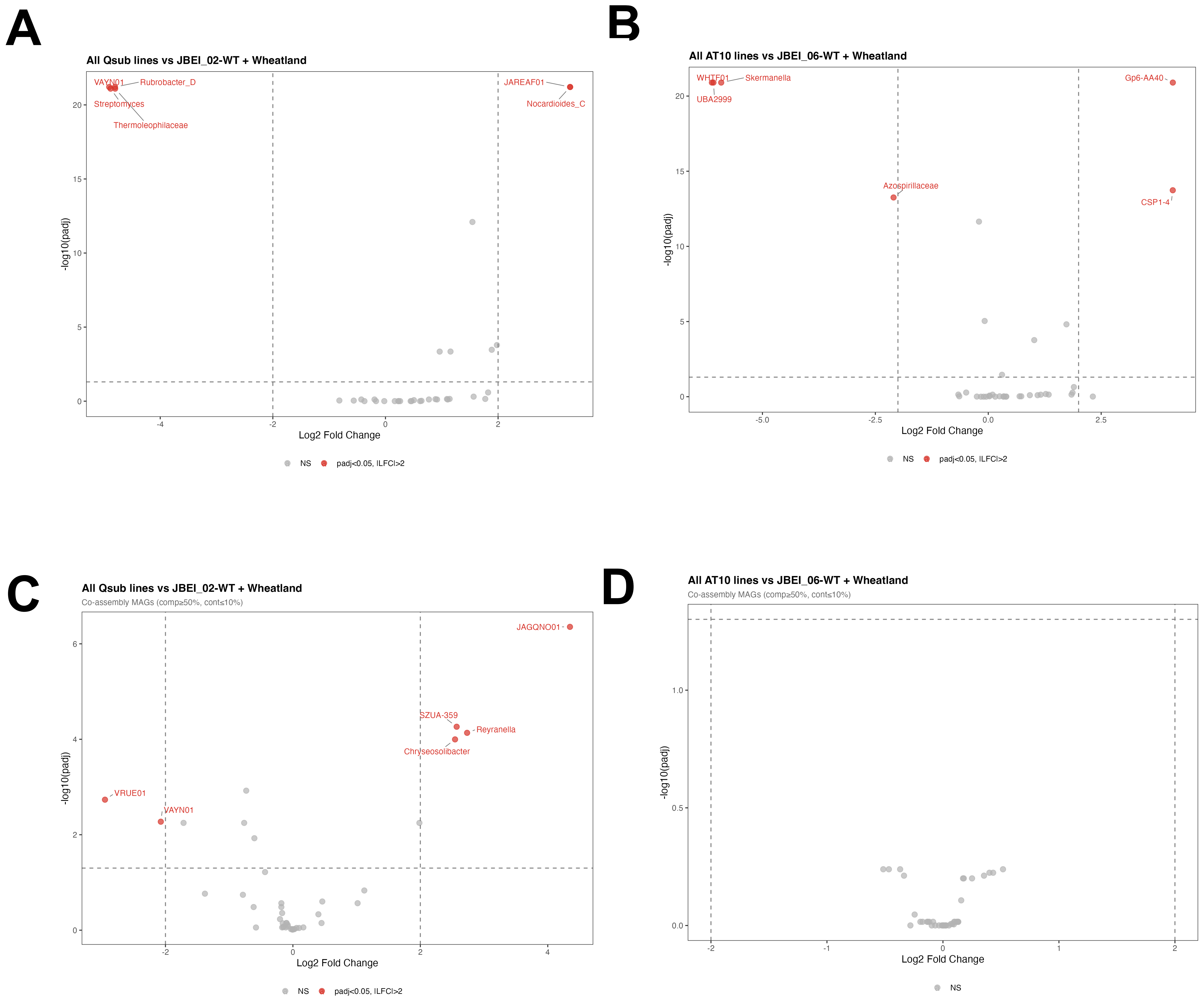


**Fig. S4. Volcano plots showing differentially abundant metagenome-assembled genomes (MAGs) identified using DESeq2.** Results are shown for the Qsub (**A**, **C**) and AT10 (**B**, **D**) lines based on individual assemblies (**A**, **B**) and co-assemblies (**C**, **D**). Only MAGs with a $\log_2$ fold change > 2 and a Benjamini–Hochberg (BH)-adjusted P value < 0.05 are labeled.
